# Maillard reaction-derived furan aldehydes act as positive allosteric modulators of the calcium-sensing receptor CaSR

**DOI:** 10.64898/2026.11.25.768246

**Authors:** Haruka Koizumi, Yusa Taniguchi, Kyoko Noda, Mina Takahashi, Yui Toyama, Yasuka Toda, Yoshiro Ishimaru, Eisaku Yamamoto, Yuta Yoshida

## Abstract

Maillard reaction products contribute to flavor formation and flavor-enhancing effects during food processing; however, the underlying molecular mechanisms remain unclear. We previously identified 5-hydroxymethylfurfural (5-HMF), a major Maillard reaction intermediate, in heated *shio-koji* lees and showed that it prolongs the umami aftertaste. We investigated its effect on the calcium-sensing receptor (CaSR), implicated in kokumi perception, and the umami taste receptors T1R1/T1R3. We found that 5-HMF elicited concentration-dependent intracellular Ca^2+^ responses in CaSR-expressing cells, but not in T1R1/T1R3-expressing cells. The CaSR responded to the structural analogs of 5-HMF, furfural, and 5-methylfurfural (5-MF), whereas furfuryl alcohol elicited no significant response, suggesting that the furan-2-carbaldehyde moiety primarily drives CaSR recognition. The CaSR-selective antagonist, NPS-2143, suppressed the response to 5-HMF and furfural. These compounds potentiated CaSR concentration responses to extracellular Ca^2+^, indicating their function as positive allosteric modulators. These findings reveal a molecular basis for kokumi perception elicited by Maillard reaction products.

## INTRODUCTION

The Maillard reaction majorly contributes to the production of complex flavors in heated foods, and Maillard reaction products are thought to improve flavor during food processing.^1^ However, the molecular mechanisms underlying these flavor-enhancing effects remain poorly understood. Kokumi has been proposed as a sensory attribute associated with enhanced mouthfulness, continuity, and flavor complexity.^2–4^ Representative kokumi substances include glutathione (GSH), originally isolated from garlic extracts, and γ-glutamyl-valyl-glycine (γEVG).^2,5^ Human sensory studies have shown that these compounds increase umami-related sensory perception and improve flavor continuity and mouthfeel.^2,5^

Studies suggest that the calcium-sensing receptor (CaSR), a class C G protein-coupled receptor (GPCR) that mediates extracellular Ca^2+^ sensing,^6^ plays a major role in kokumi perception.^4^ GSH and γEVG can activate CaSR in heterologous expression systems,^5,7^ and pharmacological inhibition of CaSR suppresses their kokumi-enhancing effects in human sensory evaluations.^5^ CaSR is also expressed in mouse taste buds, further highlighting its involvement in taste modulation.^8–10^ Studies have suggested an association between Maillard reaction products and CaSR signaling.^11–13^ Ahmad et al.^11^ reported that heating glucose with GSH enhances CaSR activity compared with non-heated controls. Wang et al.^12^ performed an *in silico* docking study and reported that two Maillard reaction products derived from L-theanine and D-galacturonic acid in tea bind to CaSR. Volatile short-chain aliphatic aldehydes, such as isovaleraldehyde, activate CaSR and improve salty and umami taste sensations in human sensory studies.^13^ Thus, tripeptides, amino acid-sugar reaction products, and volatile aroma compounds interact with CaSR.

Koji is prepared by culturing the fungus *Aspergillus oryzae* on grains, such as rice and soybeans, and is widely used to produce traditional Japanese foods, including sake, soy sauce, and miso.^14^ Shio-koji (salted koji) is a traditional Japanese fermented seasoning made by fermenting rice koji with salt and water.^14^ It is commonly used to tenderize meat as it contains enzymes such as aspergillopepsin I.^15^ Although sugars are not added during the fermentation of shio-koji, shio-koji lees contain substantial amounts of sugars and amino acids that are generated via the enzymatic degradation of rice starch and protein by *A. oryzae*, thereby acting as the substrates for the Maillard reaction. In our previous study, we reported that 5-hydroxymethylfurfural (5-HMF), a major Maillard reaction intermediate derived from hexose,^16,17^ is generated in shio-koji lees during heating.^18^ Additionally, 5-HMF was not detected in unheated shio-koji lees, indicating that it is produced via the Maillard reaction. Human sensory evaluation also showed that adding 5-HMF improves the umami aftertaste of consommé soup.^18^ 3-Furaldehyde, a positional isomer of furfural has been tentatively detected in heat-processed koji powder by GC-MS library matching.^19^

Because the CaSR mediates kokumi perception, we hypothesized that 5-HMF and furfural, major Maillard reaction intermediates in heated shio-koji lees, might also modulate CaSR activity. In the present study, we investigated the effects of 5-HMF and furfural on CaSR activity using a heterologous expression system to determine the chemical diversity of food-derived CaSR modulators and the molecular basis of flavor enhancement during food processing.

## MATERIALS AND METHODS

### Chemicals

Of note, 5-HMF (#H0269, Tokyo Chemical Industry Co., Ltd., Tokyo, Japan), furfural (#F0073, Tokyo Chemical Industry Co., Ltd.), 5-methylfurfural (5-MF; #137316, Sigma-Aldrich., St. Louis, MO, USA), furfuryl alcohol (#F0076, Tokyo Chemical Industry Co., Ltd.), L-phenylalanine (L-Phe; Fujifilm Wako Pure Chemical Corporation, Osaka, Japan), and glutathione (GSH; Fujifilm Wako Pure Chemical Corporation) were dissolved in a bath solution containing 140 mM NaCl, 5 mM KCl, 1 mM MgCl₂, 1 mM CaCl₂, 10 mM HEPES, and 10 mM glucose, with the pH adjusted to 7.4 using NaOH, prepared ar a stock solution of 200 mM and stored at −30°C. NPS-2143 (#CS-0319, ChemScene LLC, Monmouth Junction, NJ, USA) was dissolved in dimethyl sulfoxide at 100 mM and stored at − 30°C. Isovaleraldehyde (IVAH; #I0192, Tokyo Chemical Industry Co., Ltd.) was dissolved in dimethyl sulfoxide at concentrations of 100 mM and 1 M and stored at −30°C. Immediately before use, stock solutions were diluted in the bath solution.

### Human CaSR functional assay

The amino acid sequence of human CaSR was obtained from the National Center for Biotechnology Information (NCBI) database (accession number: XM_054348094.1). The open reading frame of human CaSR was synthesized and subcloned into the mammalian expression vector pcDNA3.1(+) by GeneArt Gene Synthesis (Thermo Fisher Scientific, Waltham, MA, USA). A consensus Kozak sequence (GCCACC) was added upstream of the start codon.

Human embryonic kidney 293T (HEK293T) cells (gifted by the Kawabata laboratory, Hirosaki University, Japan) were cultured in Dulbecco’s modified Eagle’s medium high glucose (Fujifilm Wako Pure Chemical Corporation) containing 10% fetal bovine serum (Biowest, Nuaillé, France), and penicillin–streptomycin solution (×100) (Fujifilm Wako Pure Chemical Corporation) at 37°C and 5% CO_2_.

HEK293T cells were transiently transfected with human CaSR/pcDNA3.1(+) or empty vector using ScreenFect A (Fujifilm Wako Pure Chemical Corporation). The cells were seeded onto a 96-well black clear-bottom plate (#265301, Thermo Fisher Scientific) coated with 0.1 mg/mL poly-D-lysine (#A3890401, Thermo Fisher Scientific). The cells were incubated for 48 h at 37°C and 5% CO_2_, and they were loaded with Fluo-4-AM using Calcium Kit II Fluo-4 (CS32, Dojindo, Kumamoto, Japan), according to the manufacturer’s protocol. After 1 h of incubation at 37°C and 5% CO_2_, the medium was replaced with a bath containing 1 mM Ca^2+^. Then, the fluorescence mobilization was detected using a multimode microplate reader (FlexStation 3, Molecular Devices, CA, USA) every 2 seconds for 90 seconds at 27°C. After 17 seconds of baseline measurements, chemical stimuli diluted in the bath solution containing 1 mM Ca^2+^ were added. To determine whether 5-HMF and furfural activate CaSR in a Ca^2+^-dependent manner, we alternatively used bath solutions containing 0.05, 0.1, and 0.3 mM Ca^2+^ without changing the other components, replacing the medium and buffer for the chemical stimuli. The activities of human CaSR were determined by ΔF_max_/F (“maximum fluorescence” − “baseline fluorescence”)/(“baseline fluorescence”).

All experiments were performed independently in triplicate (biological replicates, n= 3). For each experiment, the average value of ΔF_max_/F from 3–6 technical replicate wells was used as the representative value.

### Human umami taste receptor T1R1/T1R3 functional assay

A functional assay of the human umami taste receptor T1R1/T1R3 was performed as previously reported.^20,21^ HEK293T cells (a gift from the Matsunami Laboratory, Duke University, USA) were transiently co-transfected with expression vectors for hT1R1, hT1R3, rG15i2, and mt-apoclytin II. After 48 h of transfection, the cells were exposed to test stimuli and assayed for luminescence. The response from each well was calculated based on the area under the curve (AUC) and expressed as relative light units (RLU). All experiments were performed independently in triplicate (biological replicates, n = 3). For each experiment, the average RLU (AUC) from two technical replicate wells was used as a representative value.

### Statistical analyses

To characterize the dose-response relationships, fluorescence intensity data were fitted to a four-parameter log-logistic model, and EC_50_, E_max_, and Hill slope values were estimated (Fig. 1; Supplementary Fig. 1; Table 1). We analyzed dose-dependent responses using two-way analysis of variance (ANOVA), with cell type (CaSR or mock) and ligand concentration as the main factors (Fig. 1; Supplementary Fig. 1; Table 2). The effects and interactions of monosodium L-glutamate (MSG) and 5-HMF in human T1R1/T1R3 expressing cells were analyzed using two-way ANOVA, followed by Dunnett’s test for multiple comparisons (Fig. 2; Table 3). The effects of NPS-2143 were evaluated using a two-way ANOVA, with cell type and NPS-2143 as the main factors, followed by Tukey’s HSD test for multiple comparisons (Fig. 3; Table 4). To evaluate Ca^2+^-dependent responses to 5-HMF and furfural, we performed a two-way ANOVA with ligand treatment and extracellular Ca^2+^ concentration as the main factors (Table 5,6). When a significant interaction between ligand treatment and Ca^2+^ concentration was detected, the responses in the presence and absence of the ligand were compared at each Ca^2+^ concentration using Welch’s t-test with Bonferroni correction for multiple comparisons (Fig. 4, 5). All statistical analyses were performed using R software version 4.5.1 (R Core Team, Vienna, Austria). A *p*-value of <0.05 was considered statistically significant.

**Figure 1.**
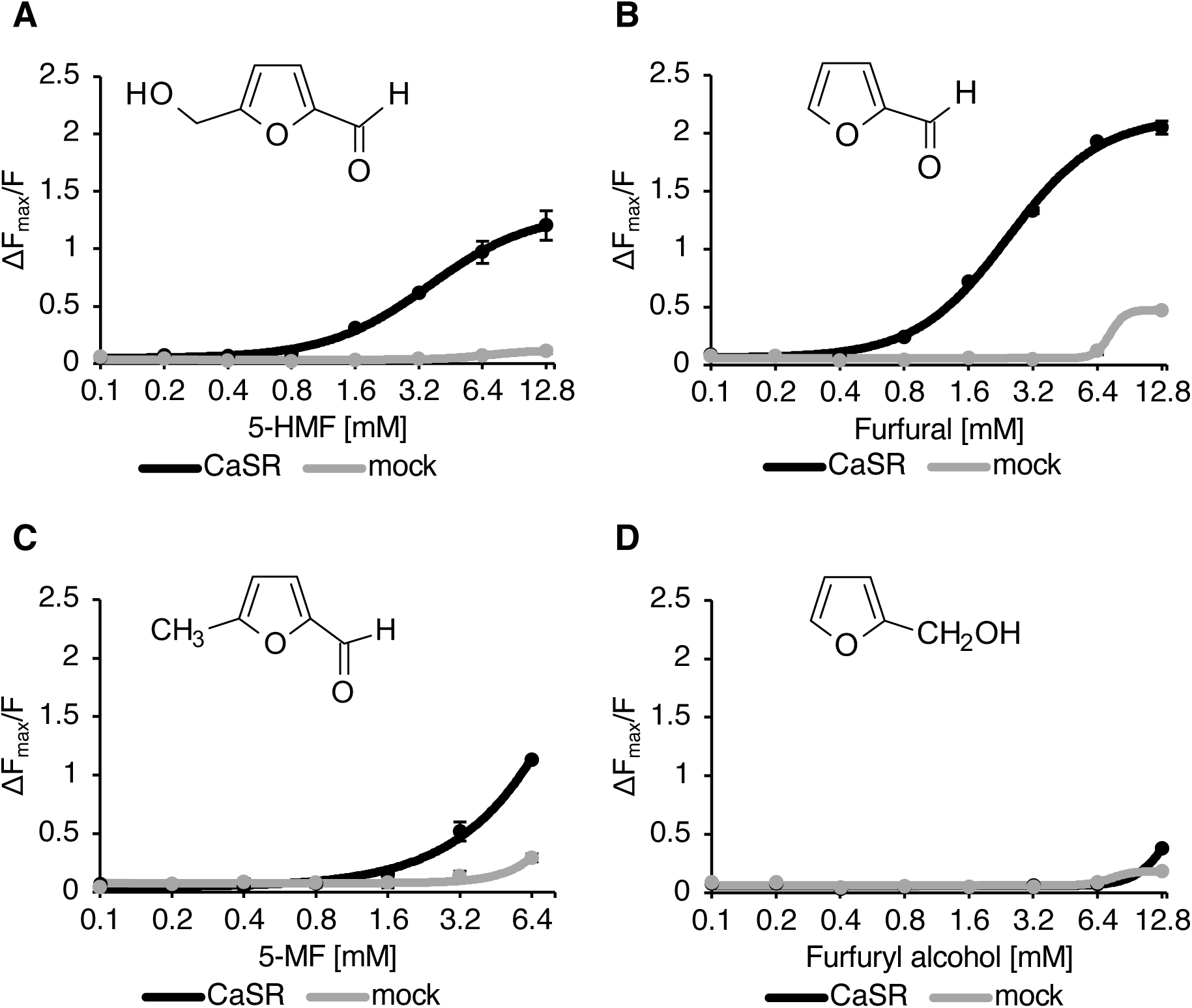
Concentration-dependent responses of CaSR to 5-HMF and structurally related analogs. (A-D) Dose-dependent relative fluorescence changes (ΔF_max_/F) in HEK293T cells expressing human CaSR (black) or transfected with mock (gray), in response to (A) 0.1–12.8 mM 5-hydroxymethylfurfural (5-HMF), (B) 0.1–12.8 mM furfural, (C) 0.1–6.4 mM 5-methylfurfural (5-MF), or (D) 0.1–12.8 mM furfuryl alcohol. Values represent means ± SEM (n = 3).

**Figure 2.**
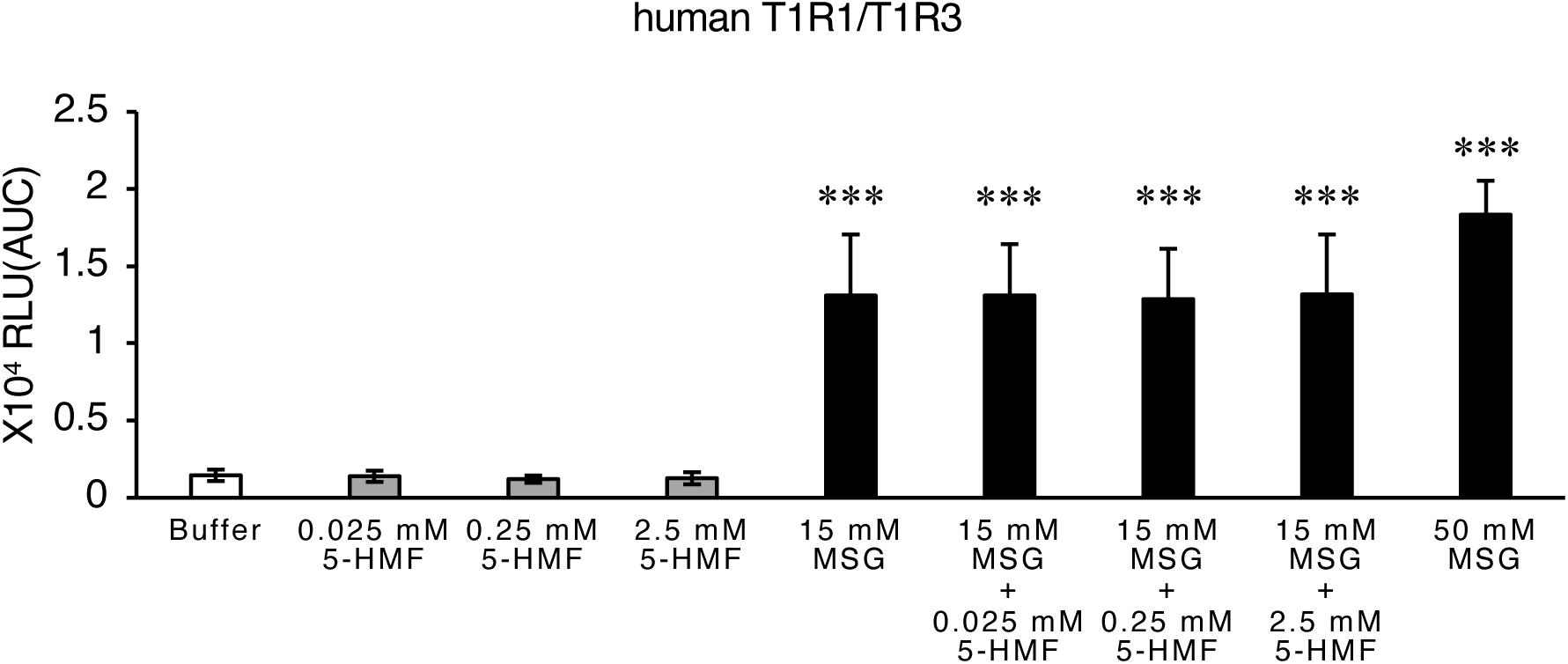
5-HMF does not modulate the human umami taste receptor T1R1-T1R3. Relative luminescence changes (RLU) in HEK293T cells expressing human T1R1/T1R3 and rat G15i2, in responses to buffer control, 0.025 mM 5-HMF, 0.25 mM 5-HMF, 2.5 mM 5-HMF, 15 mM MSG, 15 mM MSG + 0.025 mM 5-HMF, 15 mM MSG + 0.25 mM 5-HMF, 15 mM MSG + 2.5 mM 5-HMF, 50 mM MSG. \*\*\**p* <0.001 by Dunnett’s test for multiple comparisons with the buffer. Values represent means ± SEM from three independent experiments (n = 3), each performed in duplicate.

**Figure 3.**
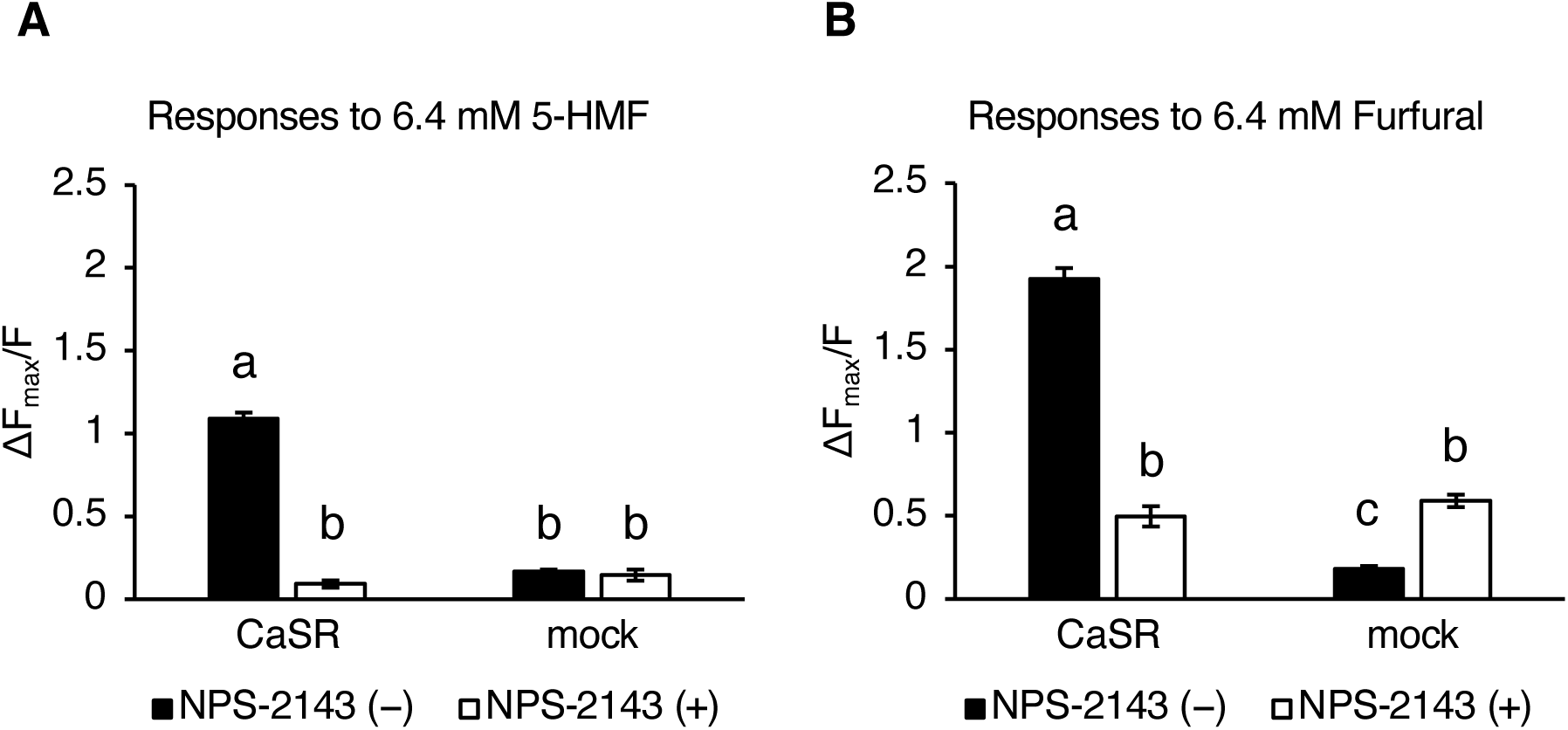
CaSR antagonist blocked the CaSR activity to 5-HMF and furfural. (A, B) Relative fluorescence changes (ΔF_max_/F) in HEK293T cells expressing human CaSR or transfected with mock, in response to (A) 6.4 mM 5-HMF or (B) 6.4 mM furfural without (black) or with (white) 10 µM CaSR antagonist, NPS-2143. Bars without a common letter are significantly different (p < 0.05; two-way ANOVA followed by Tukey’s multiple comparison test). Values represent means ± SEM (n = 3).

**Figure 4.**
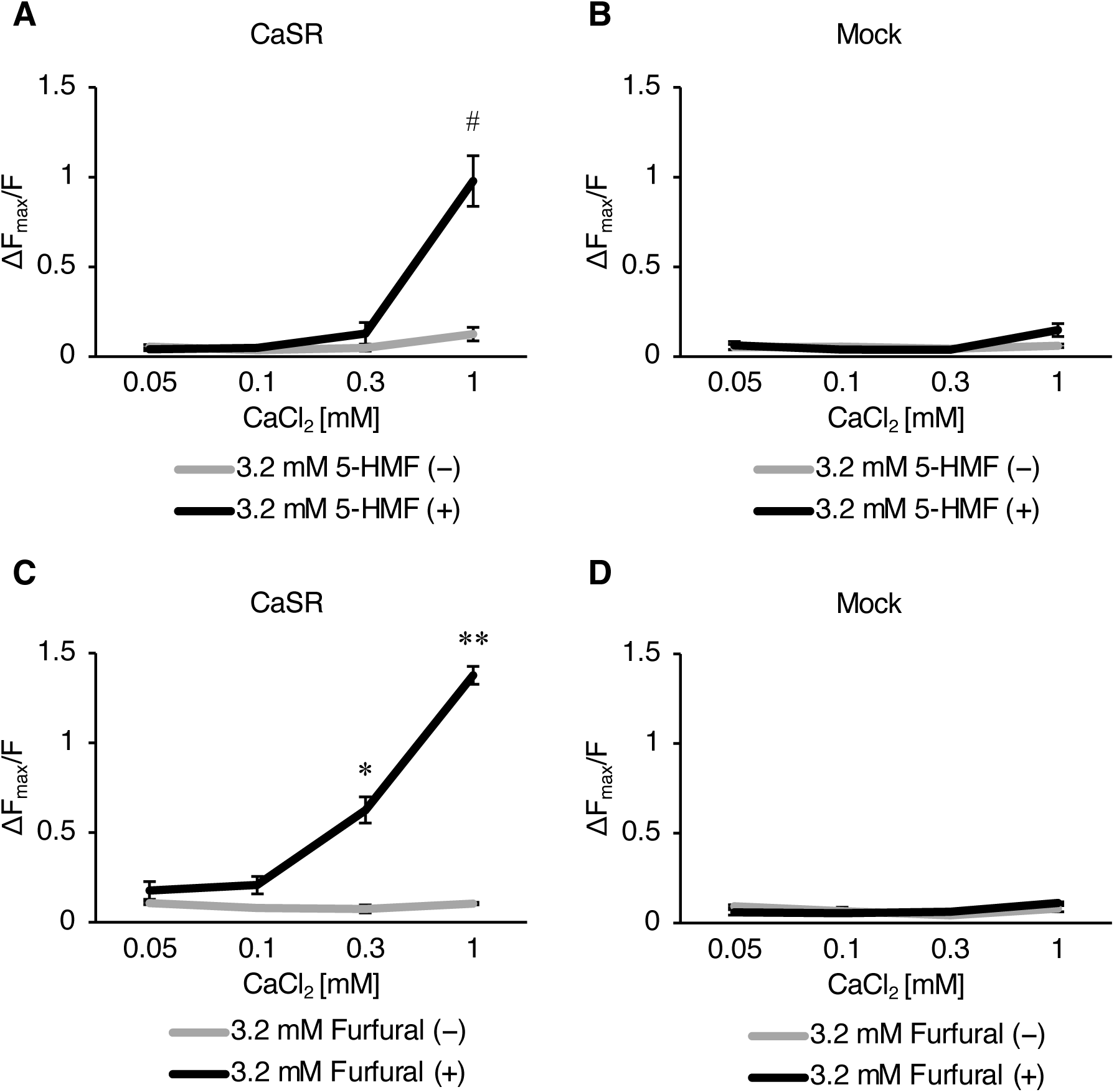
Ca^2+^-dependent modulation of CaSR by 5-HMF and furfural. (A, B) Dose-dependent relative fluorescence changes (ΔF_max_/F) in (A) HEK293T cells expressing human CaSR or (B) HEK293T cells transfected with mock, in response to 0.05, 0.1, 0.3, and 1 mM CaCl_2_ without (gray) or with (black) 3.2 mM 5-HMF. (C, D) Dose-dependent relative fluorescence changes (ΔF_max_/F) in (C) HEK293T cells expressing human CaSR or (D) HEK293T cells transfected with mock, in response to 0.05, 0.1, 0.3, and 1 mM CaCl_2_ without (gray) or with (black) 3.2 mM furfural. ^#^*p* <0.1, \**p*<0.05, and \*\**p*<0.01 by Welch’s t-test with Bonferroni correction. Values represent means ± SEM (n = 3).

**Table 1.**
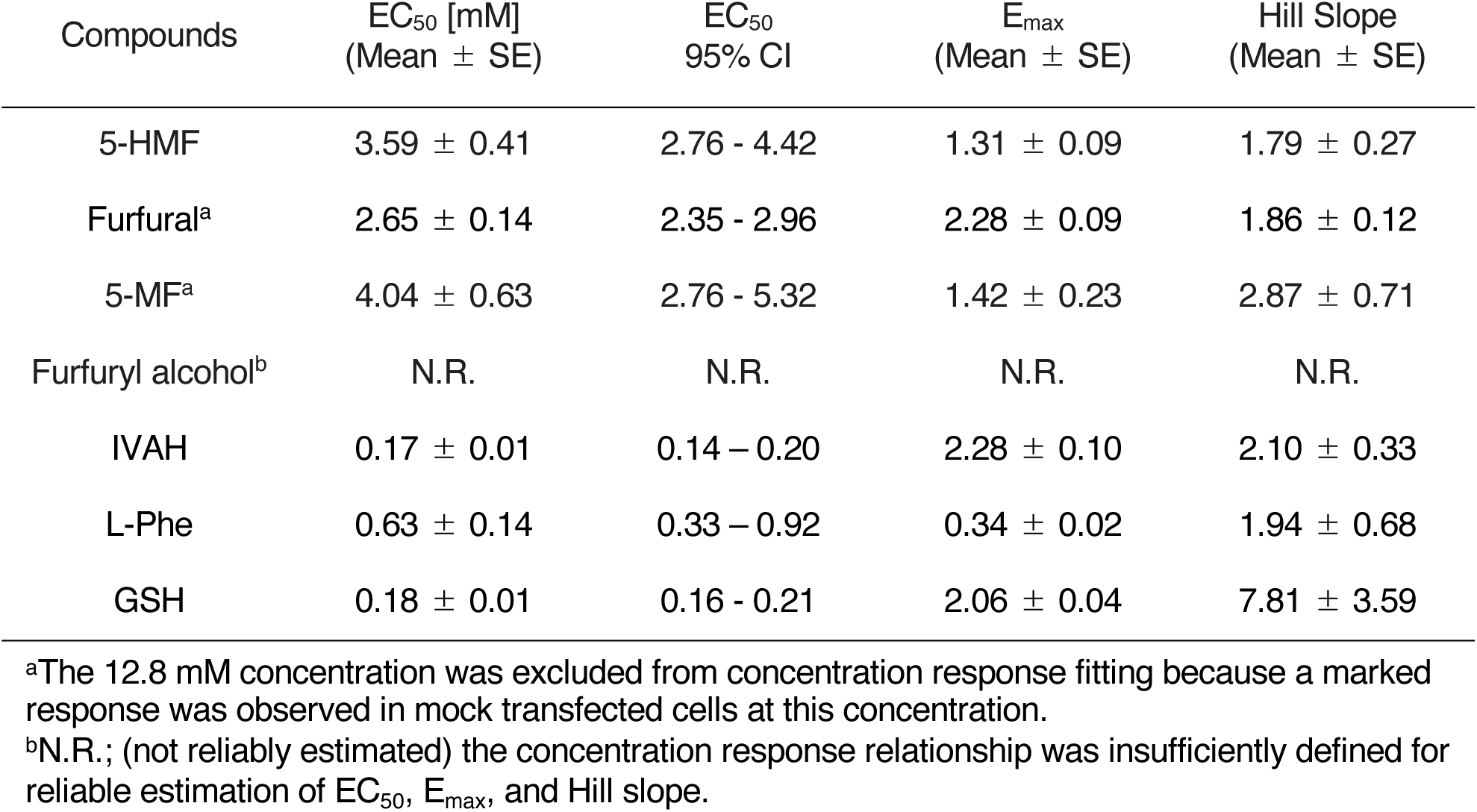
Concentration response parameters of CaSR-mediated responses to food-derived furan compounds and isovaleraldehyde (IVAH), l-phenylalanine (L-Phe), glutathione (GSH)

**Table 2.** Two-way ANOVA results of concentration responses of CaSR or mock transfected cells to food-derived furan compounds and IVAH, L-Phe, GSH.

|  | Cell type<br>(CaSR or mock) | Concentration | Cell type × Concentration |
|---|---|---|---|
| 5-HMF | *F*<sub>1,32</sub>=311.1<br>*p*<0.001 | *F*<sub>7,32</sub>=64.2<br>*p*<0.001 | *F*<sub>7,32</sub>=51.8<br>*p*<0.001 |
| Furfural | *F*<sub>1,28</sub>=4627<br>*p*<0.001 | *F*<sub>6,28</sub>=1136<br>*p*<0.001 | *F*<sub>6,28</sub>=1046<br>*p*<0.001 |
| 5-MF | *F*<sub>1,28</sub>=99.2<br>*p*<0.001 | *F*<sub>6,28</sub>=102.8<br>*p*<0.001 | *F*<sub>6,28</sub>=44.9<br>*p*<0.001 |
| Furfuryl alcohol | *F*<sub>1,32</sub>=7.8<br>*p*<0.01 | *F*<sub>7,32</sub>=52.8<br>*p*<0.001 | *F*<sub>7,32</sub>=10.4<br>*p*<0.001 |
| IVAH | *F*<sub>1,32</sub>=2843.8<br>*p*<0.001 | *F*<sub>7,32</sub>=420.0<br>*p*<0.001 | *F*<sub>7,32</sub>=387.0<br>*p*<0.001 |
| L-Phe | *F*<sub>1,32</sub>=285.9<br>*p*<0.001 | *F*<sub>7,32</sub>=15.7<br>*p*<0.001 | *F*<sub>7,32</sub>=15.5<br>*p*<0.001 |
| GSH | *F*<sub>1,32</sub>=1081.3<br>*p*<0.001 | *F*<sub>7,32</sub>=177.5<br>*p*<0.001 | *F*<sub>7,32</sub>=174.2<br>*p*<0.001 |

**Table 3.** Two-way ANOVA results of the responses of human T1R1/T1R3 transfected cells to 15 mM monosodium l-glutamate (MSG) with 0 mM, 0.025 mM, 0.25 mM, or 2.5 mM 5-HMF.

| MSG | 5-HMF | MSG × 5-HMF |
|---|---|---|
| *F*<sub>1,16</sub>=114.5<br>*p*<0.001 | *F*<sub>3,16</sub>=0.05<br>*p*=0.99 | *F*<sub>3,16</sub>=0.05<br>*p*=0.99 |

**Table 4.** Two-way ANOVA results of the responses of CaSR or mock transfected cells to food-derived furan compounds with or without NPS-2143.

|  | Cell type<br>(CaSR or mock) | NPS-2143 | Cell type ×<br>NPS-2143 |
|---|---|---|---|
| 5-HMF | *F*<sub>1,8</sub>=249.9<br>*p*<0.001 | *F*<sub>1,8</sub>=342.6<br>*p*<0.001 | *F*<sub>1,8</sub>=316.2<br>*p*<0.001 |
| Furfural | *F*<sub>1,8</sub>=269.7<br>*p*<0.001 | *F*<sub>1,8</sub>=102.2<br>*p*<0.001 | *F*<sub>1,8</sub>=333.8<br>*p*<0.001 |

**Table 5.** Two-way ANOVA results of the responses of CaSR and mock transfected cells to 3.2 mM 5-HMF or furfural under different extracellular Ca^2+^ conditions (0.05, 0.1, 0.3, and 1 mM)

|  | 5-HMF (-) or (+) | Ca<sup>2+</sup> concentration | 5-HMF × Ca<sup>2+</sup> concentration |
|---|---|---|---|
| CaSR | *F*<sub>1,16</sub>=34.1<br>*p*<0.001 | *F*<sub>3,16</sub>=38.2<br>*p*<0.001 | *F*<sub>3,16</sub>=27.1<br>*p*<0.001 |
| mock | *F*<sub>1,16</sub>=2.4<br>*p*=0.140 | *F*<sub>3,16</sub>=5.4<br>*p*<0.01 | *F*<sub>3,16</sub>=3.6<br>*p*<0.05 |
|  | Furfural (-) or (+) | Ca<sup>2+</sup> concentration | Furfural × Ca<sup>2+</sup> concentration |
| CaSR | *F*<sub>1,16</sub>=304.9<br>*p*<0.001 | *F*<sub>3,16</sub>=94.9<br>*p*<0.001 | *F*<sub>3,16</sub>=91.5<br>*p*<0.001 |
| mock | *F*<sub>1,16</sub>=0.04<br>*p*=0.845 | *F*<sub>3,16</sub>=4.4<br>*p*<0.05 | *F*<sub>3,16</sub>=2.7<br>*p*=0.077 |

## RESULTS

### 5-HMF and structurally related analogs modulate the activities of CaSR

First, we determined the responsiveness of human CaSR to 5-HMF using a heterologous expression system. Calcium imaging analyses revealed that 5-HMF increases intracellular Ca^2+^ levels in human CaSR-expressing cells in a concentration-dependent manner within the range of 0.1–12.8 mM (Fig. 1A). Conversely, 5-HMF could not induce a significant response in mock-transfected cells at any concentration (Fig. 1A).

To determine the structural characteristics associated with CaSR responsiveness, we evaluated three structurally related compounds: furfural, 5-MF, and furfuryl alcohol. Furfural induced a distinct concentration-dependent increase in intracellular Ca^2+^ levels in CaSR-expressing cells. In contrast, only a slight response was observed in mock-transfected cells at the highest concentration (Fig. 1B). Therefore, the concentration of 12.8 mM was excluded from the concentration-response analysis and statistical comparisons. Additionally, 5-MF also exhibited a concentration-dependent response in CaSR-expressing cells up to 6.4 mM (Fig. 1C). However, at 12.8 mM, 5-MF exhibited a distinct response in mock-transfected cells, reaching a level comparable to that observed in CaSR-expressing cells. Therefore, the 12.8 mM concentration was excluded from the concentration response analysis and statistical comparisons. The complete concentration response data of 5-MF, including the 12.8 mM condition, are presented in Supplementary Fig. 1A. Furthermore, 5-MF at 6.4 mM also elicited significant responses in mock-transfected cells, but a more significant response in CaSR-expressing cells (Fig. 1C). Furfuryl alcohol did not induce a clear concentration-dependent response in CaSR-expressing cells within the tested concentration range (Fig. 1D).

The responses to isovaleraldehyde (IVAH), L-Phe, and GSH, which have been previously reported as positive allosteric modulators (PAMs) of CaSR,^7,13,22^ were evaluated as positive control ligands. IVAH, L-Phe, and GSH clearly induced Ca^2+^ responses in CaSR-expressing cells (Supplementary Fig. 1B-D). The EC_50_ values for IVAH (0.17 mM) and L-Phe (0.63 mM) were lower than those previously reported for IVAH (0.681 mM^13^) and L-Phe (2.2 mM^22^), but comparable within a few-fold range, confirming the sensitivity of our functional assays (Table 1). The EC_50_ values of GSH previously reported by functional assays using CaSR-transfected HEK cells widely varied among studies, from nanomolar^5,7^ and micromolar^23^ to the millimolar range (> 3 mM by Laffitte et al.^24^; < 3 mM by Nanoff et al.^25^). The EC_50_ value of GSH found in the present study was 0.18 mM, which is within the previously reported range (Table 1).

The concentration response data were fitted to a four-parameter log-logistic model to determine the EC_50_, E_max_, and Hill slope values (Table 1). Robust concentration–response parameters were obtained for 5-HMF and furfural. For 5-MF, the model yielded estimates when concentrations reached 6.4 mM. A reliable concentration– response relationship could not be established for furfuryl alcohol. Two-way ANOVA revealed significant effects of the cell type on all four compounds (Table 2).

### The human umami taste receptor T1R1/T1R3 is not activated by 5-HMF

Umami is one of the five basic tastes, along with sweet, bitter, sour, and salty. The T1R1/T1R3 heterodimer acts as an umami taste receptor.^26,27^ Human T1R1/T1R3 mainly recognizes L-glutamate and L-aspartate^20,26^ and mediates umami synergism.^21,26,28^ Therefore, we investigated whether 5-HMF modulates human T1R1/T1R3. Two-way ANOVA revealed a significant effect of MSG on the responses of human T1R1/T1R3-expressing cells. However, 5-HMF and MSG–5-HMF showed no significant effect (Table 3). In T1R1/T1R3-expressing cells, 15 mM and 50 mM MSG induced reliable intracellular Ca^2+^ responses compared with the buffer control (Fig. 2). However, 0.025–2.5 mM 5-HMF alone could not elicit significant responses, nor did it potentiate the responses to MSG (Fig. 2). Altogether, these results suggest that 5-HMF neither could not activate or modulate human T1R1/T1R3, unlike 5′-ribonucleotides and methional.^21,28^

### Furfural and 5-HMF responses are inhibited by the CaSR antagonist NPS-2143

To further determine whether the responses to 5-HMF and furfural were mediated by the CaSR, we evaluated the effects of the CaSR antagonist NPS-2143.^29^ Treatment with NPS-2143 considerably reduced the 5-HMF-induced increase in intracellular Ca^2+^ levels in CaSR-expressing cells, whereas NPS-2143 exhibited a slight effect on mock-transfected cells (Fig. 3A). Two-way ANOVA revealed significant effects of cell type and NPS-2143 treatment and a significant interaction between these factors (Table 4).

Similarly, NPS-2143 considerably reduced the response of CaSR-expressing cells to furfural (Fig. 3B). In mock-transfected cells, furfural induced only a marginal response, consistent with the results shown in Fig. 1B. However, NPS-2143 treatment unexpectedly increased the fluorescence signal in mock-transfected cells (Fig. 3B), suggesting a CaSR-independent off-target effect of NPS-2143 rather than a furfural-specific response. Because this effect was absent under the 5-HMF condition (Fig. 3A) and occurred independently of furfural treatment, it likely reflects a non-specific action of NPS-2143 unrelated to CaSR signaling. Altogether, these results suggest that the responses induced by 5-HMF and furfural were largely mediated by CaSR.

### 5-HMF and furfural require extracellular Ca^2+^ to activate CaSR

Notably, 5-HMF and furfural can activate CaSR in a bath solution containing 1 mM Ca^2+^ (Fig. 1A, B). Based on this, we investigated whether they can also activate CaSR independently of extracellular Ca^2+^ or whether their action requires the presence of Ca^2+^, the orthosteric agonist of CaSR. We determined the responses to a fixed concentration of 5-HMF or furfural (3.2 mM) at low extracellular Ca^2+^ concentrations (0.05, 0.1, 0.3, and 1 mM). In CaSR-expressing cells, neither 5-HMF nor furfural induced a detectable response at 0.05 or 0.1 mM Ca^2+^, suggesting that these compounds do not activate CaSR in the absence of extracellular Ca^2+^ (Fig. 4A, C). As extracellular Ca^2+^ increased to 0.3 and 1 mM, both compounds elicited progressively greater responses (Fig. 4A, C). In the absence of 5-HMF or furfural, the responses remained low across all Ca^2+^ concentrations, and no comparable Ca^2+^-dependent potentiation was observed in mock-transfected cells for either compound (Fig. 4B, D).

Two-way ANOVA revealed significant effects of 5-HMF or furfural treatment on extracellular Ca^2+^ concentration, along with a significant interaction between these factors, in CaSR-expressing cells (Table 5). Welch’s t-test and the Bonferroni correction were performed for each Ca^2+^ concentration. We found that 5-HMF potentiated the CaSR response at 1 mM Ca^2+^ (Fig. 4A), whereas furfural significantly potentiated the response at both 0.3 and 1 mM Ca^2+^ (Fig. 4C). Altogether, these results indicate that the potentiating effects of 5-HMF and furfural on CaSR require sufficient extracellular Ca^2+^, supporting their role as PAMs rather than direct CaSR agonists.

### Furfural and 5-HMF potentiate the sensitivity of CaSR to extracellular Ca^2+^

To determine whether 5-HMF and furfural potentiate the sensitivity of CaSR to extracellular Ca^2+^, we measured the concentration-response relationship between CaSR and different concentrations of extracellular Ca^2+^ (1, 1.05, 1.1, 1.15, 1.2, 1.4, 1.6, and 2.4 mM CaCl_2_) in the presence or absence of 3.2 mM 5-HMF or furfural. In CaSR-expressing cells, 5-HMF and furfural increased the response to extracellular Ca^2+^ at low concentrations compared with those observed in their absence (Fig. 5A, C). No significant response to extracellular Ca^2+^ was observed in mock-transfected cells (Fig. 5B, D).

**Figure 5.**
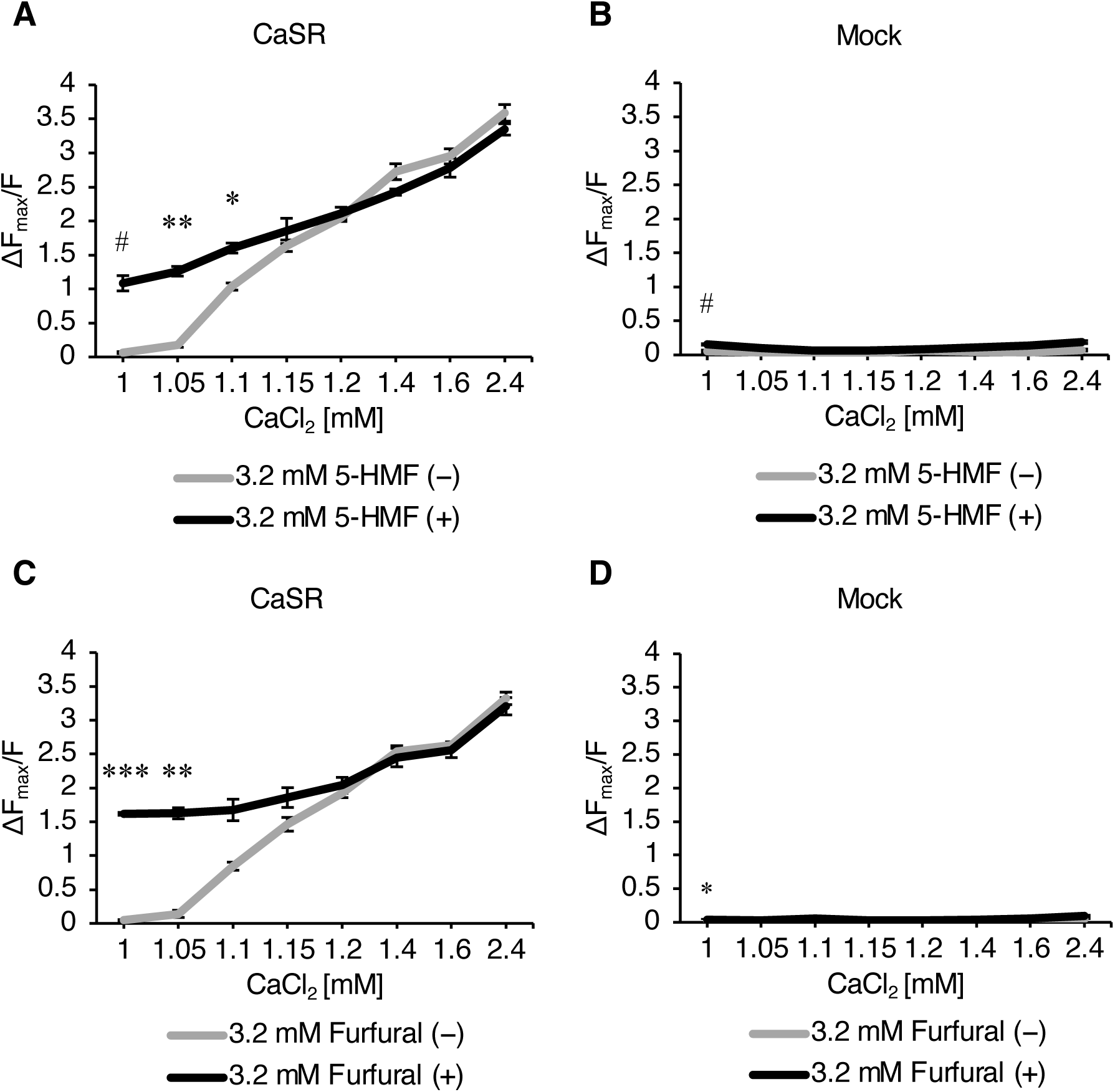
5-HMF and furfural potentiate the sensitivity of CaSR to extracellular Ca²⁺. (A, B) Dose-dependent relative fluorescence changes (ΔF_max_/F) in (A) HEK293T cells expressing human CaSR or (B) HEK293T cells transfected with mock, in response to 1, 1.05, 1.1, 1.15, 1.2, 1.4, 1.6, and 2.4 mM CaCl_2_ without (gray) or with (black) 3.2 mM 5-HMF. (C, D) Dose-dependent relative fluorescence changes (ΔF_max_/F) in (C) HEK293T cells expressing human CaSR or (D) HEK293T cells transfected with mock, in response to 1, 1.05, 1.1, 1.15, 1.2, 1.4, 1.6, and 2.4 mM CaCl_2_ without (gray) or with (black) 3.2 mM furfural. ^#^*p* <0.1, \**p* <0.05, \*\**p* <0.01, and \*\*\**p* <0.001 using Welch’s t-test with the Bonferroni correction. Values represent means ± SEM (n = 3).

Two-way ANOVA revealed significant effects of 5-HMF or furfural treatment on extracellular Ca^2+^ concentrations and a significant interaction between these factors (Table 6). Welch’s t-test and the Bonferroni correction were performed for each Ca^2+^ concentration. Furfural and 5-HMF significantly potentiated the CaSR response to lower extracellular Ca^2+^ levels (Fig. 5A, C). Altogether, these results suggest that 5-HMF and furfural potentiate the sensitivity of CaSR to extracellular Ca^2+^, providing direct evidence that they act as PAMs of CaSR.

**Table 6.**
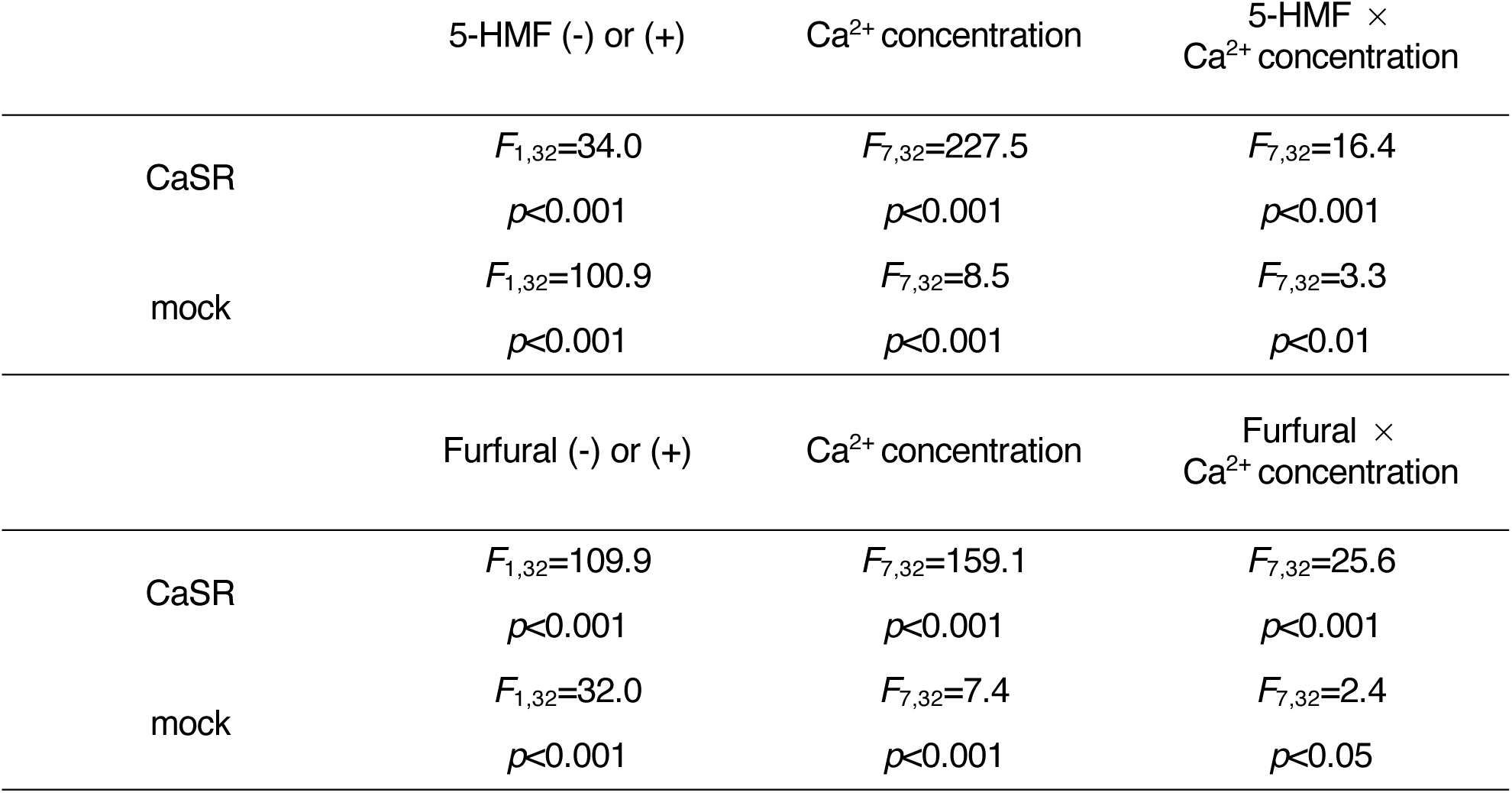
Two-way ANOVA results of the concentration responses of CaSR and mock transfected cells to extracellular Ca^2+^ (1, 1.05, 1.1, 1.15, 1.2, 1.4, 1.6, and 2.4 mM) with or without 3.2 mM 5-HMF or furfural.

|  | 5-HMF (-) or (+) | Ca<sup>2+</sup> concentration | 5-HMF × Ca<sup>2+</sup> concentration |
|---|---|---|---|
| CaSR | *F*<sub>1,32</sub>=34.0 | *F*<sub>7,32</sub>=227.5 | *F*<sub>7,32</sub>=16.4 |
|  | *p*<0.001 | *p*<0.001 | *p*<0.001 |
| mock | *F*<sub>1,32</sub>=100.9 | *F*<sub>7,32</sub>=8.5 | *F*<sub>7,32</sub>=3.3 |
|  | *p*<0.001 | *p*<0.001 | *p*<0.01 |
|  | Furfural (-) or (+) | Ca<sup>2+</sup> concentration | Furfural × Ca<sup>2+</sup> concentration |
| CaSR | *F*<sub>1,32</sub>=109.9 | *F*<sub>7,32</sub>=159.1 | *F*<sub>7,32</sub>=25.6 |
|  | *p*<0.001 | *p*<0.001 | *p*<0.001 |
| mock | *F*<sub>1,32</sub>=32.0 | *F*<sub>7,32</sub>=7.4 | *F*<sub>7,32</sub>=2.4 |
|  | *p*<0.001 | *p*<0.001 | *p*<0.05 |

## DISCUSSION

In the present study, we highlight the potential of 5-HMF and furfural—two furanic compounds formed during the thermal processing of foods—as novel PAMs of human CaSR. Structure–activity analyses of 5-HMF and its structurally related analogs (Fig. 1) showed that the furan ring and its specific substituents contribute to CaSR responsiveness. Many of our findings strongly suggest that 5-HMF and furfural act as PAMs of CaSR rather than as direct orthosteric agonists. First, the responses to both compounds were inhibited by the CaSR-selective antagonist NPS-2143 (Fig. 3), confirming the involvement of the CaSR in their activity. Additionally, both compounds required extracellular Ca^2+^ for their activity, and their potentiating effects became distinct as the extracellular Ca^2+^ approached physiological concentrations (Fig. 4). Furthermore, both compounds potentiated responses of CaSR to lower concentrations of extracellular Ca^2+^ (Fig. 5), consistent with an increase in the apparent potency of the endogenous agonist of the receptor rather than direct receptor activation in the absence of Ca^2+^. Altogether, these findings provide pharmacological evidence consistent with the classification of 5-HMF and furfural as CaSR PAMs, consistent with the functional criteria previously applied to CaSR-activating kokumi substances, including the γ-glutamyl peptide γ-EVG^5^ and, more recently, food-derived volatile compounds such as short-chain aliphatic aldehydes^13^ and medium-chain lactones.^30^ To the best of our knowledge, this is the first study to identify 5-HMF and furfural as CaSR-PAMs, thereby extending the role of CaSR in taste modulation to small furanic compounds associated with the Maillard reaction and caramelization chemistry.

Our results suggest that 5-HMF and furfural expand the structural diversity of food-derived CaSR modulators. Chemically, 5-HMF and furfural are a distinct group of Maillard reaction- and caramelization-derived furanic compounds generated during thermal processing. Notably, 5-HMF is predominantly formed from hexoses, whereas furfural is formed from pentoses.^16,31^ These compounds are widely detected in thermally processed and stored foods, including bakery products, coffee, honey, and fruit products.^32,33^ The identification of 5-HMF and furfural as CaSR PAMs adds to the growing body of evidence that chemically diverse products generated during food processing can interact with CaSR. Ahmad et al.^11^ reported that Maillard reaction products generated by heating glutathione with glucose or xylose can activate CaSR. Wang et al.^12^ reported that Maillard reaction products derived from L-theanine and D-galacturonic acid in tea can bind to CaSR, T1R1/T1R3, and OTOP1. A previous study reported CaSR-associated taste-modulating activity in thermally generated reaction products, including saltiness-enhancing peptides derived from soy protein isolate hydrolysates.^34^ Together with the present findings, these studies suggest that CaSR activity is not restricted to a single structural class of food-derived Maillard reaction products but may be affected by chemically diverse products generated during thermal processing. A recent study has also reported increased CaSR activity in reaction mixtures generated by heating meat hydrolysates with xylose, accompanied by increased levels of furfural and other Maillard-derived volatile compounds.^35^ Kim et al.^35^ reported that the individual compounds responsible for the increased CaSR activity could not be identified in complex reaction mixtures. However, their findings are compatible with the present observation that furfural can modulate CaSR activity. Further systematic examination of structurally diverse Maillard- and caramelization-derived compounds is warranted to determine how broadly this property is distributed among food-processing products. The structure–activity results obtained with 5-HMF and its close structural analogs, furfural, 5-MF, and furfuryl alcohol (Fig. 1), provide initial insights into the structural requirements for the CaSR-potentiating activity within this furanic aldehyde scaffold. Furfural, which lacks the 5-hydroxymethyl substituent of 5-HMF, can elicit substantial responses, suggesting that the hydroxymethyl group is not crucial for CaSR activity. Conversely, furfuryl alcohol, in which the aldehyde is reduced to a hydroxymethyl group, elicited no significant CaSR activity, indicating that the furan-2-carbaldehyde moiety is the principal structural determinant of the CaSR-potentiating activity. In short-chain aliphatic aldehydes reported by Kitajima et al.,^13^ the activity differed among compounds with different carbon chain lengths, whereas the present compounds retained a compact furan ring scaffold. The preservation of activity across several 5-substituted furan derivatives indicates that CaSR potentiation is not restricted to previously characterized extended aliphatic structures,^13^ and small furanic compounds associated with carbohydrate-derived thermal reactions can also exhibit CaSR-potentiating activity. We further determined whether 5-HMF could activate the human umami receptor T1R1/T1R3. However, no detectable response was observed under these test conditions. This lack of activity may be associated with the distinct chemical structure of 5-HMF compared with canonical T1R1/T1R3 ligands, which contain amino acids and 5′-ribonucleotides.^20,26^ The lack of a detectable response to 5-HMF in T1R1/T1R3-expressing cells provides an informative comparison with other food-derived taste modulators. GSH—a well-established kokumi substance and CaSR modulator—has recently been shown to act as a partial agonist and synergistic enhancer of the human T1R1/T1R3 umami receptor.^36^ Similarly, methional—a food-derived volatile aldehyde—modulates T1R1/T1R3 as a PAM and activates the CaSR.^13,21^ Thus, although GSH and methional can influence CaSR and T1R1/T1R3 signaling, 5-HMF showed detectable activity toward CaSR but not T1R1/T1R3 in the present study. These differences suggest that the receptor activity of food-derived compounds depends on their specific chemical structures and receptor context, rather than on general chemical characteristics, such as small molecular size or the presence of an aldehyde group.

The EC_50_ values obtained for 5-HMF (3.59 mM) and furfural (2.45 mM) in the present heterologous expression system were higher than those typically found in food. The concentration of 5-HMF (0.178 mM, equivalent to 22.4 mg/L) that significantly prolonged the umami aftertaste of consommé soup^18^ was approximately 20-fold lower than its CaSR EC_50_ of 5-HMF (3.59 mM, equivalent to 452.7 mg/L) obtained in the present study. However, the mechanisms underlying the discrepancy between the EC_50_ values obtained in the heterologous CaSR assay and concentration-dependent sensory effects remain unclear. Consommé soup contains multiple taste-active components, including amino acids, peptides, nucleotides, and other compounds derived from the Maillard reaction, that are absent from the defined bath solution used in the present *in vitro* assay. GSH and γEVG, which activate CaSR, are tasteless but enhance other taste sensations.^5^ Thus, the interaction of 5-HMF with other taste components in consommé soup may enhance flavor during sensory evaluation. Additionally, sensory effects *in vivo* may involve receptor systems other than the CaSR, and interactions with other taste, olfactory, or chemesthetic pathways may contribute to the lower concentrations needed for sensory effects in food. For instance, recent studies have shown that CaSR is expressed in trigeminal neurons. Additionally, γEVG enhances trigeminal responses and perceived pungency of allyl isothiocyanate, the pungent compound in wasabi.^37,38^ Similar discrepancies between flavor-enhancing concentrations and CaSR EC_50_ values have been found for other food-derived CaSR PAMs. For instance, δ-decalactone improves the fatty orosensation of pork broth at a concentration (0.18 µM) approximately 9,600-fold lower than its CaSR EC_50_ (1.73 mM^30^). Similarly, isovaleraldehyde improves salty and umami taste sensations at a concentration (0.0232 µM, equivalent to 2 ppb) approximately 29,000-fold lower than its CaSR EC_50_ (681 μM^13^).

The present study has some limitations that should be acknowledged. The EC_50_ values obtained in the *in vitro* CaSR assays varied considerably among the previous studies. For example, the reported EC_50_ values of GSH range from nanomolar to millimolar range.^5,7,24,25^ Therefore, the absolute EC_50_ values obtained from heterologous CaSR assays should be interpreted cautiously when determining the concentrations required for flavor-related effects in food. One possible explanation for the variation in the apparent EC_50_ values among the functional assays is the difference in the abundance and composition of G proteins and downstream signaling components in HEK293 cells. CaSR can activate many G protein subfamilies, including Gαq/11, Gαi/o, Gα12/13, and Gαs.^39^ HEK293 cells express a broad repertoire of endogenous G proteins and other GPCR-related signaling proteins.^40^ Therefore, differences in the endogenous complement of G proteins and downstream signaling components among expression systems may contribute to the reported variation in EC_50_ values, although this was not directly examined in this study. Additionally, the molecular binding site or the specific amino acid residues responsible for the recognition of 5-HMF and furfural by CaSR was not determined. Structural approaches, such as cryo-EM analysis and molecular docking, combined with targeted mutagenesis,^41^ would enhance our understanding of how small Maillard-associated compounds modulate receptor activity. Lastly, an unexpected response was observed in mock-transfected cells treated with furfural in the presence of NPS-2143, highlighting the possibility of non-specific effects under these experimental conditions. The basis for this observation remains unclear and warrants further investigation.

Altogether, this study demonstrates that 5-HMF and furfural—the major intermediates of the Maillard reaction—function as products of flavor formation and as PAMs of the CaSR. Our findings provide a novel receptor-based mechanism through which traditional heat-processed seasonings improve the taste and mouthfeel of food.

## Abbreviations

5-HMF,: 5-hydroxymethylfurfural
5-MF,: 5-methylfurfural
AUC,: area under the curve
ANOVA,: analysis of variance
CaSR,: calcium-sensing receptor
EC₅₀,: half-maximal effective concentration
Emax,: maximum response
GPCR,: G protein-coupled receptor
GSH,: glutathione
HEK293T,: human embryonic kidney 293T;
HEPES,: 4-(2-hydroxyethyl)-1-piperazineethanesulfonic acid
IVAH,: isovaleraldehyde; L-Phe, L-phenylalanine;
MSG,: monosodium L-glutamate
NCBI,: National Center for Biotechnology Information
PAM,: positive allosteric modulator
RLU,: relative light units
SEM,: standard error of the mean
T1R1/T1R3,: taste receptor type 1 member 1/taste receptor type 1 member 3
γEVG,: γ-glutamyl-valyl-glycine
ΔFmax/F,: relative fluorescence change.

## Acknowledgements

We thank Dr. Fuminori Kawabata (Hirosaki University) for kindly providing the HEK293T cells. We would like to thank Editage (www.editage.jp) for English language editing and their assistance in creating the graphical abstract for this paper. We thank Yuki Sato for her technical support in the initial experiments and data analysis.

## Author contributions

**Haruka Koizumi:** Data curation, Investigation, Resources. **Yusa Taniguchi:** Data curation, Investigation, Resources. **Kyoko Noda:** Writing, reviewing, and editing. **Mina Takahashi:** Investigation. **Yui Toyama:** Writing, reviewing, and editing. **Yoshiro Ishimaru:** Writing, reviewing, and editing. **Yasuka Toda:** Writing, reviewing, and editing. **Eisaku Yamamoto:** Conceptualization, Project administration, Resources, Writing, review, and editing. **Yuta Yoshida:** Conceptualization, Data curation, formal analysis, Investigation, Supervision, Validation, Writing the original draft, writing the review, and editing.

## Competing interests

This research was conducted under a joint research agreement between Ibaraki University, Ochanomizu University, Meiji University, the Institute of Science Tokyo, and Hanamaruki Foods Inc., with funding provided by Hanamaruki Foods Inc. Yui Toyama and Eisaku Yamamoto are employees of Hanamaruki Foods Inc., which manufactures heated shio-koji lees containing 5-HMF investigated in this study. The authors declare that the financial support and professional affiliation of Yui Toyama and Eisaku Yamamoto constitute potential conflicts of interest.

## Funding Sources

This study was supported by collaborative research funding from Hanamaruki Foods, Inc.

## Data availability

All raw data, information, and plasmids are available upon request.

---

**Supplementary Figure 1.**
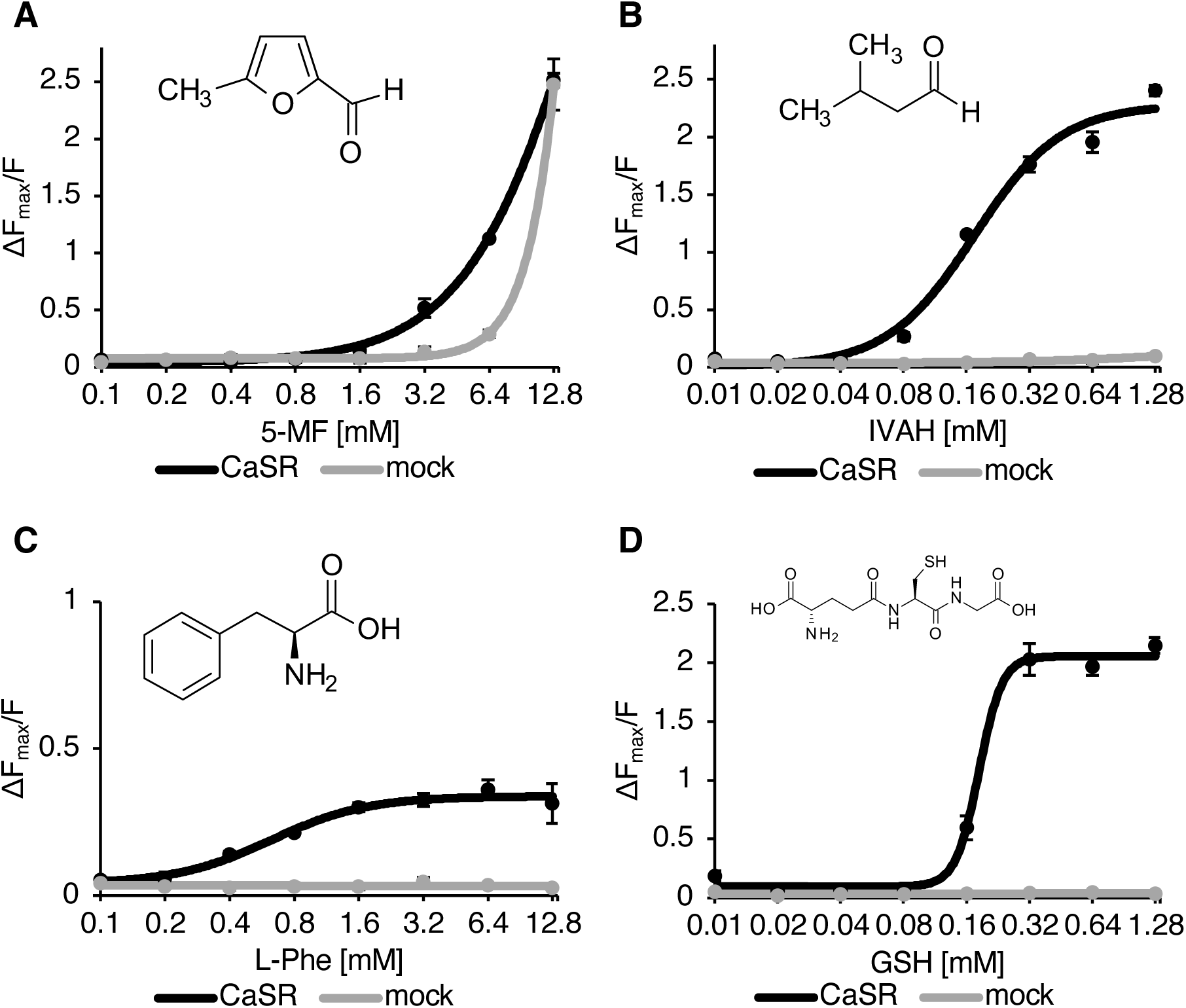
Dose-dependent activities of CaSR by 5-MF, IVAH, L-Phe, and GSH. (A-D) Dose-dependent relative fluorescence changes (ΔF_max_/F) in HEK293T cells expressing human CaSR (black) or transfected with mock (gray), in response to (A) 0.1-12.8 mM 5-methylfurfural (5-MF), (B) 0.01-1.28 mM isovaleraldehyde (IVAH), (C) 0.1-12.8 mM L-phenylalanine (L-Phe), and (D) 0.01-1.28 mM glutathione (GSH). Values represent means ± SEM (n = 3).

## Notes

### Summary of Updates

Regarding "Competing Interest Statement", we revised from "heated shio-koji lee containing 5-HMF and furfural investigated in this study" to "heated shio-koji lee containing 5-HMF investigated in this study".

